# Monitoring integrated stress response in live *Drosophila*

**DOI:** 10.1101/2023.07.13.548942

**Authors:** Peter V. Lidsky, Jing Yuan, Kseniya A. Lashkevich, Sergey E. Dmitriev, Raul Andino

## Abstract

Cells exhibit stress responses to various environmental changes. Among these responses, the integrated stress response (ISR) plays a pivotal role as a crucial stress signaling pathway. While extensive ISR research has been conducted on cultured cells, our understanding of its implications in multicellular organisms remains limited, largely due to the constraints of current techniques that hinder our ability to track and manipulate the ISR in vivo. To overcome these limitations, we have successfully developed an internal ribosome entry site (IRES)-based fluorescent reporter system. This innovative reporter enables us to label Drosophila cells, within the context of a living organism, that exhibit eIF2 phosphorylation-dependent translational shutoff – a characteristic feature of the ISR and viral infections. Through this methodology, we have unveiled tissue- and cell-specific regulation of stress response in Drosophila flies and have even been able to detect stressed tissues in vivo during virus and bacterial infections. To further validate the specificity of our reporter, we have engineered ISR-null eIF2αS50A mutant flies for stress response analysis. Our results shed light on the tremendous potential of this technique for investigating a broad range of developmental, stress, and infection-related experimental conditions. Combining the reporter tool with ISR-null mutants establishes Drosophila as an exceptionally powerful model for studying the ISR in the context of multicellular organisms.

## Introduction

The integrated stress response (ISR) is a crucial signaling pathway that allows cells to adapt to various environmental challenges and pathological conditions (Pakos-Zebrucka et al., 2016; Costa-Mattioli and Walter, 2020). This includes responses to unfolded protein stress (Walter and Ron, 2011), amino acid deficiency (Dever et al., 1992), heme depletion (Han et al., 2001), glucose deprivation (Ye et al., 2010), hypoxia (Liu et al., 2006), bacterial infections (Chakrabarti et al., 2012), and viral infections (García, Meurs, and Esteban, 2007; Stern-Ginossar et al., 2019; Sorokin et al., 2021). Research has also demonstrated the involvement of ISR in tumorigenesis (Cubillos-Ruiz, Bettigole, and Glimcher, 2017), metabolism regulation (Oyadomari et al., 2008), memory processes (Costa-Mattioli et al., 2005), development (Harding et al., 2009; Malzer et al., 2013), neurodegeneration (Bond et al., 2020), and aging (Gonskikh and Polacek, 2017; Anisimova et al., 2018). However, progress in these areas is impeded by the absence of a convenient technique for monitoring ISR in multicellular organisms.

The initiation of the ISR occurs when the translation initiation factor eIF2α gets phosphorylated at a conserved serine residue. In humans, this happens at Ser51, in drosophila at Ser50, and in C. elegans at Ser49. In mammals, the phosphorylation is carried out by specific kinases: GCN2/EIF2AK4, PERK/EIF2AK3, HRI/EIF2AK1, and PKR/EIF2AK2. Each kinase responds to a distinct subset of stress stimuli (Donnelly et al., 2013; Wek, 2018). In Drosophila, only GCN2 and PERK/PEK are present (Sood et al., 2000; Santoyo et al., 1997).

Phosphorylation of eIF2α leads to inhibition of translation for most cellular mRNA molecules due to the inactivation of Met-tRNAi Met delivery to the ribosomes. Interestingly, certain mRNAs actually increase their translation efficiency during this period (Vattem and Wek, 2004; Andreev et al., 2015; Lu, Harding, and Ron, 2004). One well-studied example of such an mRNA in higher eukaryotes is ATF4, which encodes a master transcription factor involved in cellular reprogramming to handle stress-related challenges (Wortel et al., 2017).

Cells can activate the ISR machinery as a defense response against diverse viral infections (García, Meurs, and Esteban 2007; Stern-Ginossar et al. 2019). However, viruses have evolved mechanisms to counteract this defense system (Walsh and Mohr 2011; Sorokin et al. 2021). One strategy employed by many RNA viruses involves the presence of internal ribosomal entry sites (IRESs), which initiate translation in unconventional ways (Mailliot and Martin 2018; Martinez-Salas et al. 2018; Sorokin et al. 2021). Surprisingly, certain IRESs can efficiently operate even when eIF2α is phosphorylated. In infected cells, numerous viruses have evolved specific tactics to suppress canonical translation initiation in favor of IRES-mediated protein synthesis (Walsh and Mohr 2011; Sorokin et al. 2021). A well-known example of this is the cleavage of the translation initiation factor eIF4G by the poliovirus protease 2Apro, which significantly reduces cellular protein synthesis while maintaining viral mRNA translation (Lamphear et al. 1995). Other viruses have also demonstrated similar effects through diverse mechanisms (Walsh and Mohr 2011).

IRESs are RNA elements that are able to direct eukaryotic ribosomes to internal initiation sites in a cap-independent manner (Mailliot and Martin 2018; Martinez-Salas et al. 2018; Sorokin et al. 2021). With rare exceptions, they have a highly developed secondary structure and bind some translation initiation components. Several types of IRESs and initiation mechanisms have been described, and an intergenic IRES in the cricket paralysis virus (CrPV) is one of the smallest and simplest known to date. The mechanism of action of CrPV IRES has been well-studied and it has been shown to bind to the aminoacyl site of the ribosome, mimicking the interaction between tRNA anticodon and mRNA codon during the elongation step of translation (Fernández et al. 2014; Wilson et al. 2000). As a result, the first tRNA that initiates viral polyprotein synthesis does so through an elongation-like mechanism rather than an initiation-like mechanism, and delivers alanine, not methionine (Fernández et al. 2014; Wilson et al. 2000; Martinez-Salas et al. 2018). As a result, CrPV-like IRESs do not require canonical initiation factors including eIF2 to initiate translation (Wilson et al. 2000; Pestova and Hellen 2005). Furthermore, if host mRNA translation is downregulated, CrPV-like IRESs have been shown to increase their activity (Wilson et al. 2000; Fernandez et al. 2002; Garrey et al. 2010). This stimulation is thought to be due to an increase in a number of eIF2-free 40S ribosomal subunits in the cytosol as well as relieving competition for available translation machinery. In the viral life cycle, it is believed to play a role in the switch from RNA replication to virus particle production at later stages of infection (Khong et al. 2016). The CrPV RNA contains two open reading frames: one that encodes non-structural proteins necessary for genome replication, and the other, controlled by the intergenic IRES, encoding the capsid proteins. After a sufficient number of RNA replication rounds, host mRNA translation initiation is shut down through stress-related and/or virus-derived mechanisms, resulting in a dramatic increase in IRES-mediated translation to produce enough capsid proteins for efficient virus particle assembly (Khong et al. 2016).

We have developed a fluorescent reporter utilizing a CrPV-like IRES to label stressed and virus-infected cells in live *Drosophila* organisms. Our findings demonstrate that our reporter responds to various stress stimuli known for inducing eIF2α phosphorylation and genetic manipulations that mimic ISR conditions. This reporter is particularly valuable for monitoring viral and microbial pathogenesis within living organisms. Furthermore, we generated ISR-null flies with phosphorylation-dead eIF2α^S50A^ mutation, which prevents reporter activation.

## Results

### Designing a CrPV-like IRES-based reporter

Previous research (Fernandez et al., 2002) has demonstrated the activation of CrPV-like IRES in response to stress. We verified this observation by creating a luciferase translation system directed by the CrPV IRES (Figure 1A). Following transfection of CrPV IRES-FLuc RNA in HEK293T cells, induction of oxidative stress with 100 μM arsenite, resulted in robust activation of luciferase activity (Figure 1A). The findings indicate that utilizing a reporter featuring the CrPV-like IRES offers the potential to detect cellular stress in real-time.

**Fig 1.**
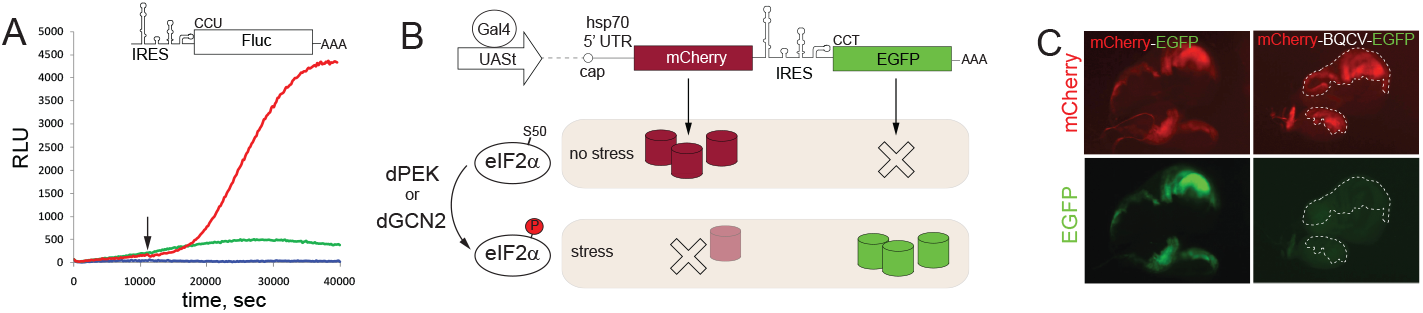
CrPV-like IRESs as stress reporters. **A**. CrPV IRES-directed translation is strongly activated in HEK293T cells after induction of oxidative stress with 100 μM arsenite. A transcript encoding firefly luciferase (Fluc) under control of CrPV IRES has been delivered by RNA transfection into the cells cultured in a plate reader and Fluc activity was continuously measured. Red line – arsenite-treated cells, green line – non-induces cells, blue line – non-transfected cells. Time of arsenite addition is shown with an arrow. **B**. A principal scheme of the reporter system construct. Under normal conditions only the cap-dependent translation of mCherry ORF is active. Upon stress, mCherry is downregulated, while the IRES-controlled EGFP-stimulated. The moderate decline in red fluorescence in stressed cells is anticipated due to slow degradation of the mCherry protein. **C**. No IRES-dependent translation might was detected in healthy wing imaginal disc. Expression of bicistronic construct or control UAS-mCherry-EGFP in-fusion construct with the use of *en-GAL4* driver demonstrated strong EGFP fluorescence. *engrailed* expression pattern was used to monitor the contrasting border between posterior and anterior compartments of the wing imaginal disc. EGFP panel in the case of the IRES-containing construct is over contrasted compared to control in-fusion construct. Posterior compartment of the wing disc is outlined with dashed lines in the former probe.

The *Drosophila* reporter comprises two components: a neutral reporter (mCherry) that is initiated by a canonical cap-dependent mechanism, marking all cells expressing the construct, and a second part (EGFP) controlled by the IRES (Figure 1B). In normal conditions, only red fluorescence should be observed, while green fluorescence appears only when cells undergo stress. The reporter was cloned into a standard vector (pUASt-attB) containing a tissue-specific UAS binding motif for expression, a minimal promoter, the 5’ untranslated region (UTR) from the Hsp70 gene, and an SV40 polyadenylation signal (Bischof et al., 2010). To serve as a control, an mCherry-EGFP fusion was also inserted into the same vector. Initially, for this study, we utilized three different IRES elements derived from CrPV, drosophila C virus (DCV) (Jousset, Bergoin, and Revet, 1977), and black queen cell virus (BQCV) (Bailey and Woods, 1977). However, careful analysis revealed that the CrPV and DCV IRESs contained cryptic polyadenylation signals within their sequences, rendering them unsuitable as stress reporters (Lidsky, Dmitriev, and Andino, 2022). Subsequently, the BQCV IRES construct was selected for further analysis, serving as the optimal choice going forward.

### BQCV IRES as a tool for stress imaging

To evaluate the utility of the BQCV-based reporter in labeling stressed cells and tissues, we conducted an initial experiment to determine background expression level. To accomplish this, the reporter and control construct were expressed in the wing imaginal discs of Drosophila larvae using the *engrailed*-Gal4 driver (as shown in Figure 1C). As expected, mCherry fluorescence was evident in the *engrailed*-positive posterior region but not in the *engrailed*-negative anterior compartment. In the case of the reporter-expressing imaginal disc, no EGFP fluorescence was detected. The sharp boundary between compartments is seen with mCherry expression. At the same time, the levels of green fluorescence between the compartments were the same, suggesting that the signal we can see is solely contributed by autofluorescence and not by reporter expression.

To assess the reporter’s activity in other tissues, we combined our construct with the pan-tissue *daughterless-Gal4* (*da-Gal4*) driver. As expected, no EGFP reporter activity was detected in larval tissues (as shown in Figure 2). Then, we used this line to test the reporter’s activation in response to stimuli that are known to induce the ISR. We exposed reporter-expressing 3^rd^ instar larvae to heat stress (37°C for 6-8 h) or an injection of 50 nl of 1M DTT. Both of these treatments are known to cause protein misfolding and activate stress responses including the ISR. The results of the exposure of the reporter-expressing flies to heat stress and DTT injection elicited tissue-specific activation of the reporter, with some unexpected results. Heat exposure mainly induced stress in the polyploid tissues of the larvae, such as the salivary glands and guts, while the injection of DTT triggered strong activation of EGFP in diploid organs, such as larval brains and imaginal discs. Therefore, our reporter indicated that ISR in *Drosophila* larvae seems to have strong tissue-specificity that also depends on the type of induction.

**Fig 2.**
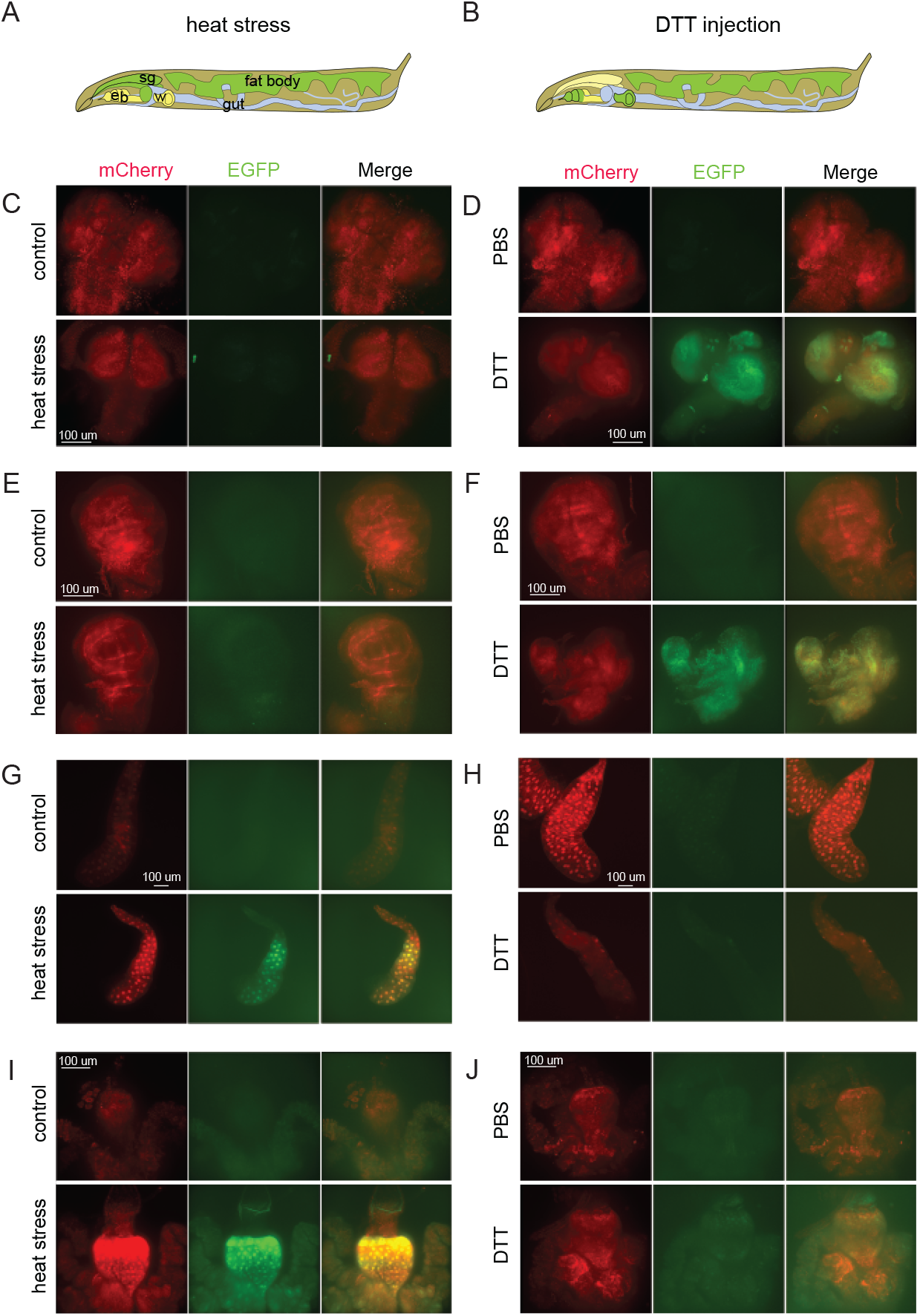
Differential patterns of stressed tissues induced by heat and DTT injection. 3^rd^ instar *Drosophila* larvae with two copies of reporter construct and two copies of *da-GAL4* driver were exposed to prolonged heat stress (6-8 h, 37°C; **A, C, E, G, I**) or injected with 50 nl of 1M DTT (**B, D, F, H, J**). Heat stress was not able to induce reporter activation in the brain tissue (**C**) and imaginal discs (**E**), but activated strong EGFP signals are the polyploid larval salivary glands (**G**) and in the midguts (**I**). On the contrary, DTT injection did not induce reporter activation in larval tissues (**H, J**) but activated reporter in the brains (**D**) and imaginal discs (**F**). The results are summarized on schemes (A, B). All imaging conditions are equivalent within organ groups.

### BQCV IRES activity depends on eIF2α^S50^ phosphorylation as judged by genetic analysis

Subsequently, we employed *Drosophila* genetics to modulate the ISR and evaluate the regulatory influence of eIF2αS50-P on the activation of our reporter. We used an active mutant of GCN2 kinase (Bjordal et al. 2014). In addition, we constructed a phosphorylation-dead eIF2α^S50A^ knock-in mutant (Fig 3A) and UAS-driven mutant variants of eIF2α (Fig 3B).

**Fig 3.**
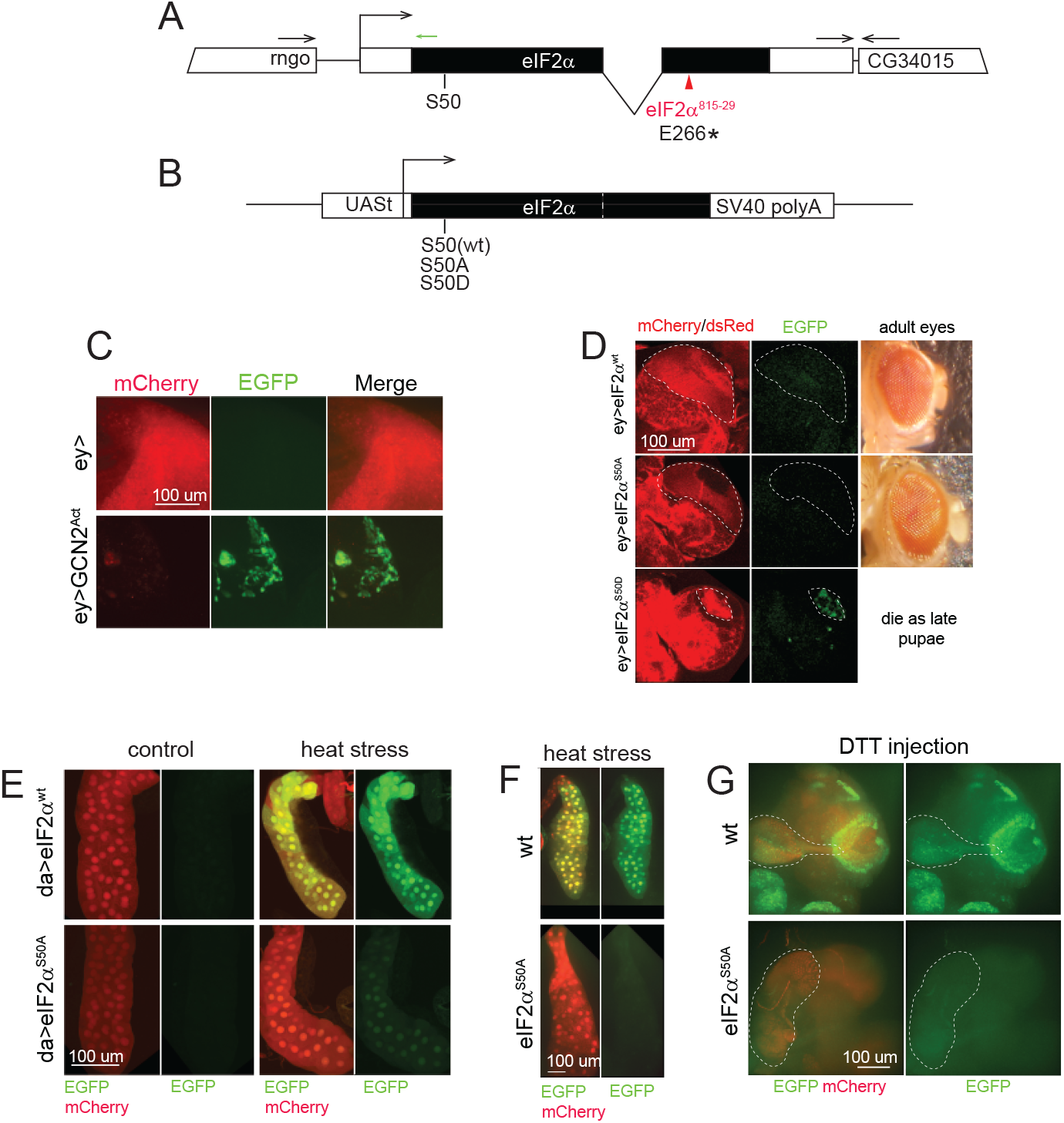
Genetic manipulations with ISR machinery affect BQCV IRES-based reporter activation. **A**. Structure of *eIF2α* gene. Gene spans are shown as rectangles, Coding sequences are in black. Location of *eIF2α*^*815-29*^ lesion was mapped as amber mutation E266* and shown in red. Annealing region of the CRISPR gRNA used for S50A edit is shown as a small green arrow. **B**. Scheme of the constructs for UAS-mediated expression of eIF2α variants. **C**. Co-expression of the reporter and GCN2 constitutively active mutant in the optic lobes of the third instar larvae brain induces a decrease in mCherry levels and an increase in EGFP levels. Note also a decline in the size of the optic lobe due to the toxicity of the construct. **D**. Co-expression of eIF2α variants in the optic lobes as in panel **C**. Phosphorylation-mimicking eIF2α^S50D^ variant induces reporter activation, while eIF2α^wt^ and phosphorylation-dead eIF2α^S50A^ do not. Optic lobes are encircled with dashed lines. Red fluorescence outside of these areas comes from 3xP3-dsRed marker eIF2α variants are labelled with. **E**. Co-expression of eIF2α^S50A^ but not eIF2α^wt^ reduces the stress reporter activation in response to heat (7 h, 37°C). Expression of the reporter and eIF2α variants in this experiment was mediated by *da-GAL4* as in panel **2G. F, G**. *eIF2α*^*S50A*^ mutant obtained with a CRISPR editing cannot mount stress reporter activation in response to heat (**F**) or DTT injection (**G**) applied as in **Fig 2**. All imaging conditions are equivalent within the panels.

First, we expressed the reporter in the optic lobes of *Drosophila* larval brains in combination with a constitutively active mutant of the GCN2 kinase that constantly phosphorylates eIF2α and induces ISR (Figure 3?). Control brains showed bright mCherry signals without any signs of EGFP fluorescence. As expected, expression of mutant GCN2 resulted in a strong reduction of optic lobes sizes and diminished the mCherry expression. Strong EGFP signals were detected upon expression of the mutant GCN2 kinase.

Next, we have generated constructs encoding the eIF2α variants under control of the UAS promoter (Figure 3B). The wild-type eIF2α, a phosphorylation-dead mutant S50A, and a phosphomimetic mutant S50D were engineered similarly to the described constructs but with use of UAS instead of heat-shock promoter (Qu et al. 1997). The transgenic proteins were shown to be fully functional since the wild-type and the S50A mutant were capable to partially rescue the known *eIF2α*^*815-29*^ lesion (McKim, Dahmus, and Hawley 1996), which we identified as an amber mutation in position 266 of eIF2α (Figure 3A). Rescued animals were strongly underrepresented among wild-type progeny, however displayed no apparent deleterious phenotype (Fig S1). UAS/*ey*-Gal4-mediated ectopic expression of the phosphorylation-mimicking eIF2α^S50D^ mutant results in lethality at pupal stages. We have analyzed the effect of eIF2α mutants expression on stress reporter activation in the presence of the endogenous wild type gene. As expected, expression of the phosphomimetic mutant eIF2α^S50D^ but not eIF2α^S50A^ and eIF2α^wt^ in the optic lobes induced reporter activation (Figure 3D). Conversely, a *da*-Gal4-driven ubiquitious expression of a phosphorylation-dead eIF2α^S50A^ mutant resulted in a dramatic reduction in EGFP expression upon the heat stress (Figure 3E). The inhibition was incomplete perhaps because of the activity of wild type eIF2α produced with the genomic copy. Phosphorylated eIF2α can lock GTP exchange factor eIF2B in inhibited state (Kenner et al. 2019; Gordiyenko, Llácer, and Ramakrishnan 2019), thus, if a portion of eIF2α is phosphorylated, this might enough to suppresses translation considerably (Siekierka, Manne, and Ochoa 1984). This explains our observation that the induction of endogenous eIF2α phosphorylation partially inhibited translation even in presence of non-phosphorylated eIF2α.

In these experiments, we found that the BQCV IRES-based stress reporter significantly relies on the phosphorylation status of eIF2α at Ser50. This observation highlights the crucial role of eIF2α Ser50-phosphorylation in regulating the expression of stress reporter.

### Engineering CRISPR edited *eIF2α*^*S50A*^ *Drosophila* mutant

Next, we produced a mutant fly with the S50A mutation introduced into *eIF2α* genomic copy. In brief, guide RNAs and Cas9-encoding plasmid were co-injected into fly embryos together with a donor construct encoding eIF2 with the S50A substitution, mutation in PAM-site sequence, and a *3xP3-dsRed* selection marker. dsRed-positive flies were collected, balanced, and the mutations were validated by Sanger sequencing. *eIF2α*^*S50A*^ hemizygous males and homozygous females were viable and fertile, and were kept as a stock. We have combined S50A allele with BQCV-based reporter and *da-Gal4* driver insertion. Mutant flies exposed to heat stress (Figure 3F) and DTT injection (Figure 3G) demonstrated no EGFP reporter activation. Thus, eIF2α phosphorylation plays a critical role for activation of the BQCV IRES-mediated translation at least in case of these stress inducers.

### Imaging of stressed tissues in *Drosophila* larvae

To further demonstrate the utility of our imaging technique, we exposed reporter-expressing animals to heat stress (Figure 4A), viral (Figures 4B-D), and bacterial infections (Figures 4E-F). As before, larvae exposed to heat (Figure 4A) displayed strong reporter activation in polyploid tissues (see also Figure 2). Infection with DCV by feeding led to strong activation of reporter in a subset of animals (Figure 4B). We detected reporter activity in multiple tissues of EGFP-positive individuals including salivary glands (Figure 4C) and imaginal discs (Figure 4D). Brain tissue remained EGFP-negative until the late stages of infection (Figure 4D). Infection with a bacterial pathogen, *Pseudomonas entomophila*, is known to induce ISR in the gut of infected animals (Chakrabarti et al. 2012). Consistently, feeding larvae with *P. entomophila*, but not with a non-pathogenic strain of *Escherichia coli*, demonstrated a bright fluorescence in the gut visible *in vivo* (Figure 4E) and in the dissected organs (Figure 4F). Amino acid starvation in 2^nd^ instar larvae also resulted in activation of EGFP expression in the midgut (Figure 4G). We concluded that our reporter system is robustly responding to stress stimuli and can be used for visualizing stressed cells *in vivo* and in dissected tissues.

**Fig 4.**
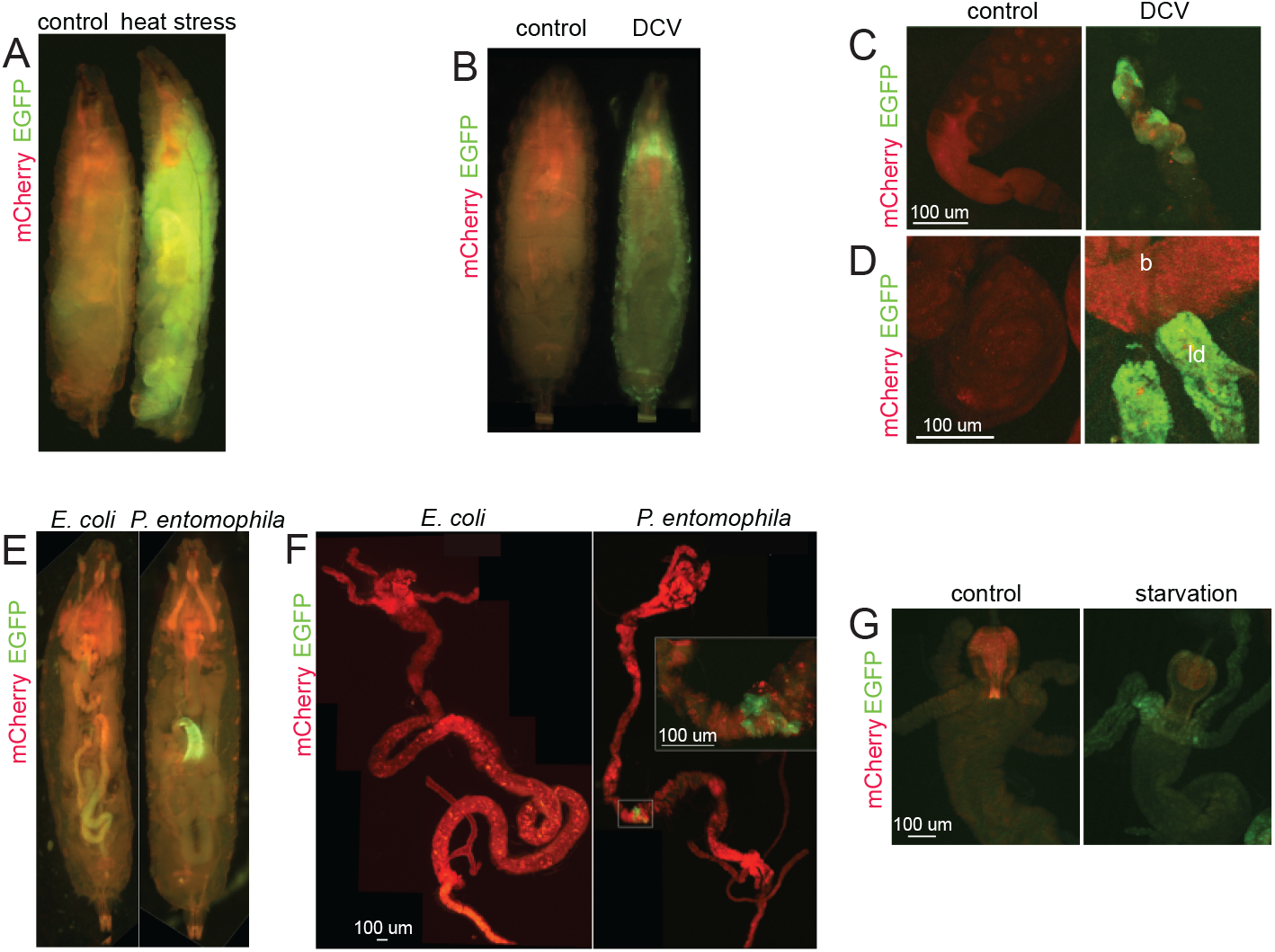
BQCV IRES-based reporter activation in response to heat, infection, and starvation. Homozygous flies expressing two copies of reporter constructs and two copies of *da-GAL4* were used in all experiments. **A**. 3^rd^ instar Drosophila larvae were exposed to heat as in **Fig 2**, and subjected to full body imaging. **B**. Larvae infected with DCV occasionally express strong reporter activation in multiple tissues including salivary glands (**C**) and imaginal discs (leg discs are labeled as ‘ld’ in panel **D**). Brains (‘b’ in **D**) remain EGFP-negative in these animals. **E**. Feeding larvae with *P. entomophila* but not *E. coli* induces reporter activation in the gut. **F**. Localization of the stressed cells within the gut confirmed by dissection of the infected animals. **G**. Amino acid starvation of 2^nd^ instar larvae induces stress reporter activation in the guts. All imaging conditions are equivalent within the panels.

## Discussion

### Reporter for imaging stress *in vivo*

The Integrated Stress Response (ISR) plays a pivotal role in numerous pathological conditions. Its functions go beyond cellular adaptation to environmental challenges like chemical intoxication, starvation, and infectious diseases. Mounting evidence highlights the involvement of ISR in diverse areas, including development (Pakos-Zebrucka et al., 2016), brain function (Sharma et al., 2020; Zhu et al., 2019), aging (Derisbourg, Hartman, and Denzel, 2021; Derisbourg et al., 2021), metabolic regulation (Harding et al., 2003), and various other pathologies (Costa-Mattioli and Walter, 2020). While extensively studied in cultured cells, the investigation of ISR in multicellular organisms remains largely unexplored. This paper introduces novel tools for studying ISR in living *Drosophila* animals.

CrPV-type IRESs are activated when the host’s translation is halted due to certain conditions (Khong et al., 2016; Garrey et al., 2010; Wilson et al., 2000; Fernandez et al., 2002). In this study, we developed a BQCV IRES-based reporter system in Drosophila to visualize stressed cells and tissues under various damaging conditions (Figures 2 and 4). This innovative system allows labeling of cells experiencing ISR-induced host translational shut-off in vivo (Figure 4) and enables direct monitoring of these cells in living organisms without the need for dissection, fixation and additional staining, as previously required for ATF4-based stress reporters (Kang et al., 2015).

Our analysis indicated that different tissues are differentially susceptible to different stressors: polyploid larval tissues were more susceptible to heat, while the diploid imaginal and brain tissues – to the DTT-mediated induction (Figure 2). These observations suggest the existence of tissue-specific variations in the induction of translational stress responses that could be explained by differential sensitivities of the tissues to ISR induction as well as variable basal protein synthesis and metabolic activities, as well as concentration of translation machinery components and codon adaptation indices (see Anisimova et al. 2023; Allen et al. 2022 for a review).

### Tools for genetic manipulation of the integrated stress response

To confirm the dependence of BQCV-IRES mediated translation on ISR and expand the *Drosophila* toolkit for stress investigation, we have developed UAS-based constructs. These constructs encode wild type eIF2α, as well as phosphorylation-dead S50A and phosphomimetic S50D mutants (Qu et al. 1997). This enables targeted manipulation of the ISR in specific tissues, as shown in Figures 3D-E.

Expression of constitutively active GCN2 kinase, as well as eIF2α^S50D^ results in induction of BQCV IRES-mediated EGFP expression (Figures 3C-D). Conversely, the presence of eIF2α^S50A^ mutant reduces the reporter activation induced by heat stress (Figure 3E). An entire removal of wt *eIF2α* allele by introduction of S50A site into the endogenous chromosome by means of CRISPR editing resulted in a complete inhibition of stress-induced activation of BQCV IRES-based reporter (Figures 3F-G) indicating that ISR plays a critical role in reporter induction.

Importantly, *Drosophila* animals are able to survive to adulthood and stay fertile having the non-phosphorylatable eIF2α^S50A^ variant as the only source of eIF2 activity, indicating ISR is unessential for development in this model. eIF2α phosphorylation is essential for mouse development: homozygous phosphorylation-dead mutants are dying 18 h after birth because of defective gluconeogenesis (Scheuner et al. 2001). Conversely, *eIF2α*^*S49A*^ mutants in *C. elegans* have been reported to be viable (Rollins, Lind, and Rogers 2017). Now we extend this observation to the *Drosophila* model. Therefore, phosphorylation-dead *eIF2α* mutant invertebrates provide an opportunity to investigate the role and regulation of ISR in the context of the living organism.

### Imaging of infection-induced stress *in vivo*

Virus and bacterial infections are variable and complex processes. Our knowledge of virus infection within the organism is still quite limited. An accurate imaging of stressed cells in the context of infection *in vivo* might provide critical information for the understanding of pathogen-host interplay.

Most experiments on fluorescent labeling of virus-infected cells have been done with recombinant viruses, expressing fluorescent proteins (Costantini and Snapp 2015) or smaller tags (Sakin et al. 2016). The major limitations here are the molecular constraints of virus genomes, attenuating effects of the fluorescent inserts, and rapid evolution of viruses that tend to discard the tag (Mueller and Wimmer 1998). Some reporter systems have been designed to track virus infections with the constructs encoded in cell genome (see e.g. Emmott, Sweeney, and Goodfellow 2008; Hwang, Chen, and Yates 2006), however, these approaches are specific for a cleavages performed by virus-specific proteases and only rarely go beyond the tissue culture (Ekström and Hultmark 2016). The ISR or alternative types of inhibition of host translation are hallmarks of infection with many different pathogens and therefore our reporter system is expected to be useful for imaging various infections.

We expect that our techniques will push forward the ISR research in the Drosophila model and will be specifically useful for study of pathogen-host interactions in invertebrates.

## Materials and Methods

### Plasmids

Stress reporters. To construct *pUASt-mCherry-BQCV_ires-EGFP-attB* plasmid, encoding the IRES-based stress reporter, *pUASt-attB* vector was opened with EcoRI and XbaI enzymes. Fragments encoding EGFP and mCherry were amplified by PCR (see Table SI for oligonucleotide sequences). EGFP fragment was digested with BglII and XbaI. mCherry fragment was treated with EroRI and BsrGI. IRES sequences were synthesized by Genescript and excised from their vector plasmid with BsrGI and BglII. *pUASt-EBFP2-BQCV_ires-mCherry-attB* was generated essentially as *pUASt-mCherry-BQCV_ires-EGFP-attB* with use of the corresponding fluorescent protein sequences.

*pUASt-eIF2α*^wt^*-attB, pUASt-eIF2α*^*S50A*^*-attB* and *pUASt-eIF2α*^*S50D*^*-attB* were produced by introduction of the corresponding fragments into *pUASt-attB* backbone, opened with EcoRI/XbaI enzyme pair. The mutagenized gene fragments were synthesized by Genescript, digested with EcoRI/BamHI restriction enzymes and introduced into the plasmid along with an invariant part of the gene, prepared with BamHI/XbaI enzymes.

*pScarlessHD-DsRed_eIF2α_S50A_PAM* donor construct for CRSIPR editing was constructed by insertion of mutagenized version of eIF2α sequence into *pScarlessHD-DsRed* (gift of Kate O’Connor-Giles (Brown University, Providence, RI), Addgene plasmid #64703) sequence with SapI and AarI restriction sites. The resulting construct harbored S50A mutation and a GCC->GCT synonymous transition in A31 inactivating the PAM site. *pU6-BbsI-chiRNA_eIF2α* gRNA *eIF2α*-targeting sequence (GTACTCAAGCAGATGAACGT) was cloned into *pU6-BbsI-chiRNA* (Gratz et al. 2013) opened with BbsI.

Plasmid encoding firefly luciferase under control of CrPV IRES was described earlier (Prokhorova et al. 2016).

### mRNA transfection with continuous monitoring of luciferase activity

Polyadenylated CrPV-Fluc mRNA was *in vitro* transcribed using a 50T-tailed PCR product as a template, as described previously (Prokhorova et al. 2016). Human HEK293T cells (ATCC) were seeded in DMEM supplemented with 10% FBS onto a 96-well white opaque FB plate (Greiner). 12-16 h later, the cells were transfected with 30 ng of mRNA per well using GenJector-U (Molecta, Russia). Beetle luciferin (Promega) was added to the medium to final concentration of 0.5 mM, and the plate was immediately placed into Infinite 200 Pro microplate reader (TECAN) provided with a gas control module. The cells were incubated at 36.6°? in 5% CO_2_ for 12 h, luciferase activity was continuously measured every 5 min with integration time 5 s, as describes earlier (Panova et al. 2023).

### Flylines and fly husbandry

Reporter and eIF2α variants-encoding attB constructs were inserted into M-ZH-86Fb.attP docking site (Bischof et al. 2010). Classic *en-GAL4* insertion was used (Yoffe et al. 1995). *GAL4-da*.*G32* (Wodarz et al. 1995) insertion was on the third chromosome and was driving Gal4 ubiquitously; in the text, it is referenced as *da-Gal4. UAS-GCN2*^*Act*^ insertion (Bjordal et al. 2014) was generously provided by Prof. Pierre Leopold (University of Nice). *y, w, ey-Flp, Act5c>CD2>Gal4* (Pignoni and Zipursky 1997) chromosome was ascribed in the text as *ey>Gal4* and was kindly provided by Prof. Hugo Stocker (ETH, Zurich).

Fly stocks were kept at 18°C, while the crosses and infected flies - at 25°C. Cleaning the stocks from potentially contaminating viruses was performed by bleaching the eggs. Anti-*Wolbachia* treatment was performed by supplementing the fly food with tetracycline. Screening for the presence of *Wolbachia* was performed by PCR.

Feeding flies with pathogens was performed by adding them to the food.

Injection larvae with 50 nl of 1M DTT in PBS was performed using the Drummond Scientific Nanoject II nanoinjector.

### Pathogens

Drosophila C virus sample was a kind gift of Prof. Jean-Luc Imler (University of Strasbourg) and was amplified in Drosophila Schneider’s cells. *Pseudomonas entomophila* sample was a gift of Prof. Bruno Lemaitre (EPFL, Lausanne).

### Organ samples

Dissected organs were fixed with 4% PFA for 20 min, rinsed with PBS, quenched with 1M glycine on PBS for 10 min, washed with PBS and mounted on Vectashield (Cole-Parmer) containing the DAPI stain.

### Microscopy and image analysis

Whole larvae images were obtained with Nikon SMZ1500 and Leica MZ16 F. Images were deconvolved with Helicon Focus software (HeliconSoft). Acquisition of confocal images stacks was performed with a Leica TCS SP2 confocal microscope equipped with 10x and 40x objectives, with an Olympus FV1000 confocal microscope equipped with 10x and 60x objectives, Nikon Ti Spinning Disc confocal microscope equipped with 20x and 40x objectives, with Nikon FN-1 microscope equipped with 10x, 20x, and 40x lenses, and with Nikon Crest LFOV Spinning Disk/ C2 Confocal equipped with 10x, 20x, and 40x lenses. Images were processed using Imaris software (Bitplane), Fiji (Schindelin et al. 2012) or NIS-Elements (Nikon).

## Acknowledgements

The work was supported by the Impetus Longevity Grant and NIH R01AI137471 for R.A. and by the Russian Science Foundation grant no. 18-14-00291 to S.E.D.

## Figure legends

**Fig S1. Validation of UAS-eIF2α constructs. A**. Gene structure of eIF2α. The green arrows correspond to the oligos, and the line – to the PCR product that was used to genotype *eIF2α*^*815-29*^ (E266*) mutation. Both oligos anneal in intronic sequences. Thus, the intronless UAS-eIF2α constructs (**B**) are not recognized by this oligo pair. **C**. Adult males rescued with UAS-eIF2α^wt^ and UAS-eIF2α^S50A^ driven with *da*-Gal4 driver.

