## Supplementary figures and images for "Monitoring integrated stress response in live *Drosophila*"

### Fig S1

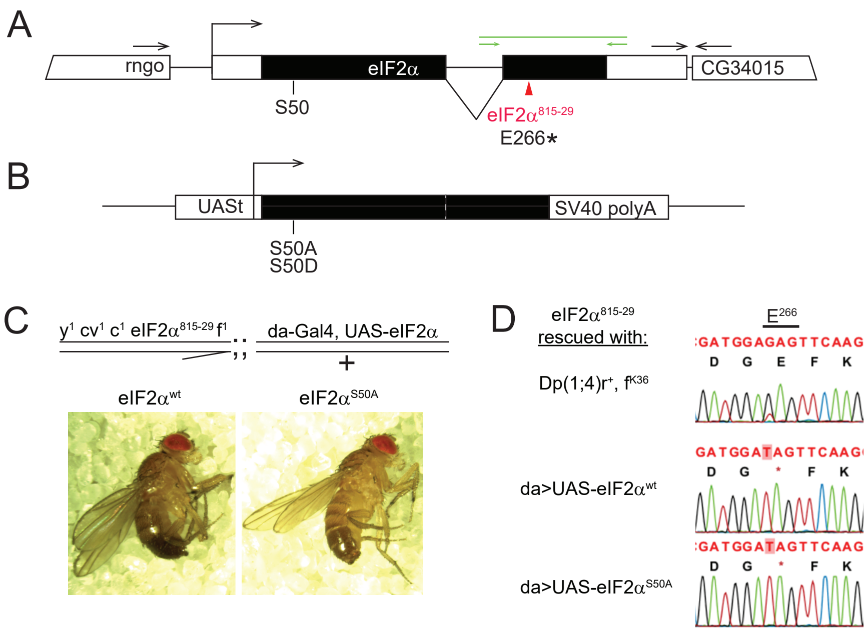
